# An optimal-fitness framework for modeling perceptual compression

**DOI:** 10.1101/2023.02.23.529655

**Authors:** Victor Quintanar-Zilinskas

## Abstract

Perceptual systems are constrained by their information transmission capacity. Accordingly, organismal strategies for compressing environmental information have been the subject of considerable study. The efficient coding model posits maximized mutual information between stimuli and their neural representation. The reward maximization model posits minimized signal distortion, operationalized as reward foregone due to stimulus confusion. The matched filters model posits the preferential transmission of information that informs evolutionarily important decisions. Unfortunately, the efficient coding model is sometimes at odds with empirical findings, and all three models struggle with recapitulating each other’s predictions. Here I aim to reconcile the models by developing a framework for modeling compression in which: compression strategies dictate stimulus representations, compressed stimulus representations inform decisions, decisions deliver rewards, environments differ in decision-reward associations and fitness function, and therefore, different environments select for different compression strategies. Using this framework, I construct environments in which the fittest compression strategy: optimizes signal distortion, optimizes both signal distortion and mutual information, and optimizes neither but nevertheless is fit because it facilitates the avoidance of catastrophically risky decisions. Thus, by modeling compression as optimal with respect to fitness, I enable the matched filters model to recapitulate the predictions of the others. Moreover, these results clarify that mutual information maximization and signal distortion minimization are favored by selection only under certain conditions. Hence, the efficient coding model is reconciled with the findings that it fails to predict, because those findings can now be understood to derive from outside the model’s proper scope of application. Going forward, the optimal-fitness framework is poised to be a useful tool for further developing our understanding of nature’s perceptual compressions; a salient reason why is that it enables empirical findings to be bridged not only with concepts from information theory, but also economics.

**Author Summary:** Perceptual systems are constrained by their information transmission capacity. Thus, stimuli are not transmitted in full detail, but are instead compressed. Presently, there are several extant models of compression that are supported by empirical results. However, they do not recapitulate each other’s predictions, and are not bound by any common conceptual framework. In the present study, I create a common conceptual framework: the optimal-fitness framework, which allows for the evaluation of the evolutionary fitness of a particular compression in a particular environmental context. This framework, in turn, allows me to define the features of the environments that favor the compressions predicted by the extant models. These findings serve to refine the extant models by defining their domain of applicability, and to unify the models by demonstrating the existence of environments in which their predictions overlap. Furthermore, the optimal fitness framework accommodates the expression of, and the demonstration of the evolutionary value of, various naturalistically plausible compressions that are not predicted by the existing models.

## Introduction

Perceptual systems inform their host organism’s decisions, but their ability to do so is constrained by their information transmission capacity. Accordingly, within the field of perceptual science, numerous models have been proposed to describe how perceptual systems compress their environment’s information. Typically, these models propose a compression strategy, predict the stimulus and environmental state distinctions that it enables, and offer an account for why natural selection would favor its use.

The efficient coding model (1, 2) is perhaps the exemplar of a well-studied and empirically-corroborated perceptual compression model. Formally, it posits the maximization of mutual information (MI) between internal perceptual representations and environmental states; stated alternatively, if a perceptual system can only transmit *β* bits, it will transmit those that maximize discrimination amongst the environment’s states. This model is corroborated by findings that the input-output curves of individual sensory neurons are optimally adapted to encode naturalistic stimulus intensity distributions (1). This model is further corroborated by findings that neuron populations in visual, auditory, and olfactory systems exhibit: information-maximizing features (such as reduction of redundancy between neurons) that manifest specifically during the processing of naturalistic stimuli (3-7), stimulus coding that is adapted to the distribution of naturalistic stimuli (8), and code plasticity when the distribution of their environment’s stimuli changes (9, 10). Also, MI maximization is an evolutionarily plausible compression strategy because it limits the surprise associated with an organism’s action-driven state transitions, and thereby facilitates the organism’s pursuit of high-value states (11).

Alternative models of compression posit that perceptual systems preferentially transmit information that informs evolutionarily-important decisions, while tolerating the confounding of environment states that motivate similar actions. When no further details are specified, this compression strategy characterization is known as the matched filters model (12), because the information filtered from the environment matches the organism’s needs. Empirically, it appears to be validated by moth auditory neurons that are maximally sensitive to the sound frequencies used by bat echolocation (13, 14), by moth auditory processing that extracts stimulus features that inform escape behavior (15, 16), and by the existence, in several other species, of various other perceptual adaptations seemingly specialized for the processing of stimuli that inform essential survival actions (17-20). A more specific relationship between compression and decision-making is proposed by the reward maximization model: it predicts that compressions are describable by rate-distortion theory, and that the distortion objective minimized is total stimulus confusion cost, because minimizing confusion cost is equivalent to maximizing the arithmetic mean reward (AMR) delivered by an organism’s decisions. This model is appealing both because of the intuitive relationship between AMR and evolutionary success (21, 22) and because it accurately predicts stimulus discrimination performance in many experimental settings (21, 23).

Unfortunately, because it has not been demonstrated that these models recapitulate each other’s predictions, and because the evolutionary advantages conferred by the proposed compression strategies are by no means well-characterized, there is presently no model that provides a unified understanding of perceptual compression. Here, I describe several limitations of the extant models. The efficient coding model is difficult to reconcile with a significant body of empirical findings. For example, it does not predict reward-driven stimulus discrimination plasticity, which enables organisms to recognize reward-linked stimuli with heightened specificity (21, 24-27). It also does not intuitively account for why, in numerous moth species, all auditory neurons are tuned to the same frequency (13); while this configuration equips moths to distinguish between “bat present” and “bat absent” environmental states, it seems ill-suited for maximizing distinctions across environmental states in general. Evolutionarily, the implication of these deviations from the model’s predictions is that nature foregoes MI-maximizing compressions in favor of those that support the pursuit of organismal objectives. The reward maximization model, meanwhile, is difficult to reconcile with empirical findings because the nature-imposed costs of experimentally-observed perceptual errors are often unknown to experimenters. If stimulus confusion costs are unknown, it cannot be claimed that the observed compression minimizes their incurrence. Evolutionarily, the limitation of the reward maximization model is that natural selection sometimes favors the pursuit of organismal objectives that deviate from AMR maximization (28). In such cases, one would expect selection to favor compressions aligned with these alternative objectives. Finally, while the matched filters model’s evolutionary premise is difficult to refute, its predictive power is limited to species for which the major decision challenges (e.g., predator evasion) are well-characterized. For species that navigate a rich diversity of decision scenarios (e.g., most mammals) and process a wide range of stimuli, this model, unlike the models premised on either MI or AMR maximization, presently provides no specific basis for predicting which stimuli will be confounded.

In the present study, with the aim of extending the matched filters model and bridging it with the efficient coding and reward maximization models, I develop a mathematical framework that expresses the evolutionary favorability of compressions that inform adaptive decisions. Within this framework, environments are defined by the stimuli they present, the stimuli-informed decisions they require, and their fitness function. Also, within each environment, organisms make decisions that are informed by compressed stimulus representations, and these representations are determined by the organism’s compression strategy. Thus, an organism’s fitness-determining decision outcomes are a function of its compression strategy. Thus, optimizing fitness reveals the compression strategy favored by natural selection in a particular environment.

Using this framework, I explore which compression strategies are favored in several different environments, and obtain the following results. First, I find that there are environments in which AMR-maximizing compressions are optimally fit. This result achieves the integration of the reward maximization model into the optimal-fitness framework. Second, I find that there are environments in which AMR is maximized but MI is not, in which AMR is maximized while MI is well-approximated, and in which both AMR and MI are maximized. These results show that the efficient coding model can also be integrated into the optimal-fitness framework. Also, the environments that are most favorable to MI-maximizing compressions exhibit selection pressures similar to those experienced by many vertebrates, which is notable because efficient coding has frequently been observed in vertebrate perceptual systems. That said, these validations of the fitness sometimes conferred by efficient coding are complemented by results that confirm the long-held suspicion (1) that MI maximization is sometimes at odds with fitness maximization. Thus, efficient coding is not a generally-applicable model of perceptual compression. Third, I find that there are environments in which AMR-maximizing compressions are outcompeted by compressions optimized for goals such as risk avoidance, opportunistic resource pursuit, and minimized variability of resource acquisition performance. Thus, reward maximization is not a generally-applicable model of perceptual compression either. Instead, this study’s findings can collectively be regarded as a theoretical delineation of the settings in which the efficient coding and reward maximization models are appropriate to invoke. Outside of these settings, the optimal-fitness framework is very well-suited for expressing the diversity of decision-making challenges to which real-world perceptual systems are adapted.

## Results

### Details of the optimal-fitness framework

As stated above: environments present organisms with stimuli, organisms make decisions informed by compressed representations of the stimuli, and decision outcomes determine fitness. More specifically: stimuli, compressed representations, fitness-determining decisions, and environments are formulated as follows. Stimuli take the form of items, which are defined by a set of quantitative attributes. The possible values of individual attributes can span either a radial or linear range, and thereby exist within the interval [−*π, π*] or [0, *b*_*a*, 1_] (subscript *a* differentiates the linear-range attributes), respectively. Together, the attribute axes define an environment’s “item space”. Compressed representations are necessitated by perceptual systems that are able to transmit only *β* bits about each item; in lieu of transmitting attribute values, perceptual systems map items to one of *m* (*m* = 2^*β*^) percepts. These item-percept mappings, which are determined by item attribute values and the parameterization of the organism’s compression strategy, are a lossy compression operation: multiple items are mapped to the same percept, possible real-world items outnumber the possible percepts, and attribute value information is not fully preserved (Table 1). Fitness-determining decisions present organisms with a decision scenario objective and *n* items sampled randomly from the environment, and require the choice of one item. Here, fitness is a monotonically increasing function of the sum of the reward values gained from the items chosen over an organism’s lifetime of decisions. Also, item reward values are determined by item attributes and are scenario-specific (to illustrate: different item attributes are rewarding to an organism afflicted by hunger vs. by thirst). Accordingly, organisms exploit the item attribute information conveyed by item-percept mappings by favoring some percepts over others during item selection decisions, and furthermore by exhibiting scenario-specific percept preferences. Finally, environments are characterized by their fitness function, their probability distribution of items within item space, their ensemble of decision scenarios, and the transmission capacity (*β*) of their inhabitant organisms. In some instances, biologically meaningful insight can be derived by specifying only some environment properties, and subsequently finding the compression strategy that is optimal for the entire set of environments that share the specified properties.

**Table 1.** A simple mapping of real-world items to internal percepts.

| Item | 1 | 2 | 3 | 4 | 5 | 6 |
| --- | --- | --- | --- | --- | --- | --- |
| Attribute 1 | 1 | 2 | 3 | 4 | 5 | 6 |
| Attribute 2 | 7 | 8 | 6 | 5 | 7 | 5 |
| Percept | 0 | 0 | 0 | 1 | 1 | 1 |

This 1-bit perceptual system maps items to Percept 1 if their Attribute 1 value exceeds 3.5. Otherwise, it maps them to Percept 0.

Two elements of this framework that have also been used in previous studies are the conceptualization of compression as the mapping of a multitude of stimuli onto a much smaller number of percepts (29-31), and the evaluation of compression strategies on the basis of total reward gained while repeatedly choosing one item from amongst *n* (31, 32).

In order to facilitate the derivation of insights applicable to real-world perceptual systems, many of the present study’s environments are designed with features that mimic those of natural environments, that simplify the derivation of the optimal compression strategy, or perhaps do both. One of these features is naturalistic geometry. Conceptually, naturalistic geometry is the notion that items with similar attributes often have similar scenario-specific reward values and often are mapped to the same percept. More formally, in this study’s environments with naturalistic geometry: scenario-specific item reward values are determined by a single attribute; individual scenarios (denoted by subscript *j*) are specified by the reward-determining attribute, the reward-maximizing attribute value, and the reward conferred by items with that attribute value (*M_j_*); and item value decreases linearly with distance from the reward-maximizing attribute value at a rate that is uniform across all scenarios. Three additional environment features, which in the present study’s environments with naturalistic geometry can be assumed unless otherwise specified, are that *n* is constant across decision scenarios, decision scenarios are equally probable, and item probabilities are uniform throughout item space. Finally, several of this study’s environments are, with respect to organismal decision-making, unidimensional, which is to say that the reward-determining attribute is shared across all decision scenarios. Conceptually, optimizing compression strategies in unidimensional environments is useful, both because the theorized optimal compression can be evaluated against the many published experiments that have measured discrimination amongst stimuli that vary along a single attribute, and because the biological implications of the theorized optimal compression are often straightforwardly discernable. Within the optimal-fitness framework, optimizing compression strategies in unidimensional environments is convenient, because it reduces to the placement of percept boundaries along the span of the reward-determining attribute. To illustrate: when the reward-determining attribute’s values span a linear range, the attribute’s maximum value can be labeled *b*_1_ (rather than *b*_*a*,1_), percept boundaries *b_i_* can be indexed in order of proximity to *b*_1_, percepts *P_i_* can be indexed according to their upper boundary, and thus, fitness can be written as a function of the *b_i_* (*i* = 2: *m*; *b*_1_ is a constant and *b*_*m*+1_ = 0).

### Selection for maximization of arithmetic mean reward

Although environments that favor an AMR-maximizing compression strategy are somewhat of a theoretical contrivance, I nevertheless elaborate their properties, and those of the favored compressions, in order to advance understanding of naturalistic perceptual compressions that are well-described by the reward maximization model. For brevity, I refer to this set of environments as E1. In E1, only the following properties are specified: item presentation probabilities and scenario-specific item reward values are presumed constant through evolutionary time, and fitness is linearly proportional to AMR. In some naturalistic settings, it is plausible that an organism’s contribution to the next generation’s gene pool would be roughly proportional to its total reward accumulated over a very long lifetime of decisions.

Under these conditions, the compressions that maximize AMR also minimize signal distortion; specifically, the total cost of stimulus confusion. For organisms with noiseless item-percept mapping, I verify the previously-intuited (22) equivalence of these objectives in Parts 1.1 and 1.2 of S1 Appendix. Next, I consider AMR maximization in the context of a more realistic (33, 34) perceptual process model, in which both encoding (item-percept mapping) and decoding (item identity inference, based on encoded percept) are noisy. These noise sources can result in an item’s attributes being estimated in accordance with the attributes of a non-target percept (an item’s target percept is the percept to which it’s mapped in noise’s absence). I show in Part 1.3 of S1 Appendix that AMR is maximized by compression strategies that compensate for noise, henceforth referred to as noise-adjusted optimal mappings.

Further understanding of AMR-maximizing compression strategies can be developed by optimizing compression in environments within a subset of E1 that exhibits naturalistic geometry, E2, and more specifically within E2A, E2B, E2C, and E2D, all of which are unidimensional subsets of E2. For simplicity, item-percept mapping within E2 is assumed to be noiseless unless otherwise specified, and *M_j_* is assumed to be uniform across all decision scenarios and written simply as *M*.

In E2A, there is only one decision scenario, item reward value is maximized at *b*_1_, and the environments within E2A are thus differentiated only by *n* and *β*. Because of E2A’s simplicity, I am able to present, in Part 2.1 of S1 Appendix, a closed-form expression for the fitness-maximizing *b_i_* within all of its constituent environments. Then, in Part 2.2 of S1 Appendix, I use the E2A environment in which *n* = 3 and *β* = 1 to illustrate the difference between noiseless-case and noise-adjusted optimal mappings.

### Selection for maximization of mutual information

In the previous section, the optimal-fitness framework is shown to incorporate the reward maximization model of compression. This is achieved by constructing framework environments that select for compression strategies that conform to the model’s predictions. Here, I explore the extension of this approach to the efficient coding model.

I begin this endeavor by examining, in E2A_β1_ and E2B, the fitness conferred by MI-maximizing compression strategies. E2A_β1_ is the subset of E2A environments in which *β* = 1. E2B is similar to E2A in that there is only one decision scenario in which item reward value is maximized at *b*_1_, but differs as follows: *β* and *n* are respectively fixed at 1 and 2, and item probabilities across the span of the value-determining attribute are distributed as illustrated in Fig 1A. Note that the environments in E2B are differentiated only by their value of *a*, and that the environment in which *a* = 0 is also a constituent of E2A_β1_.

**Fig 1.**
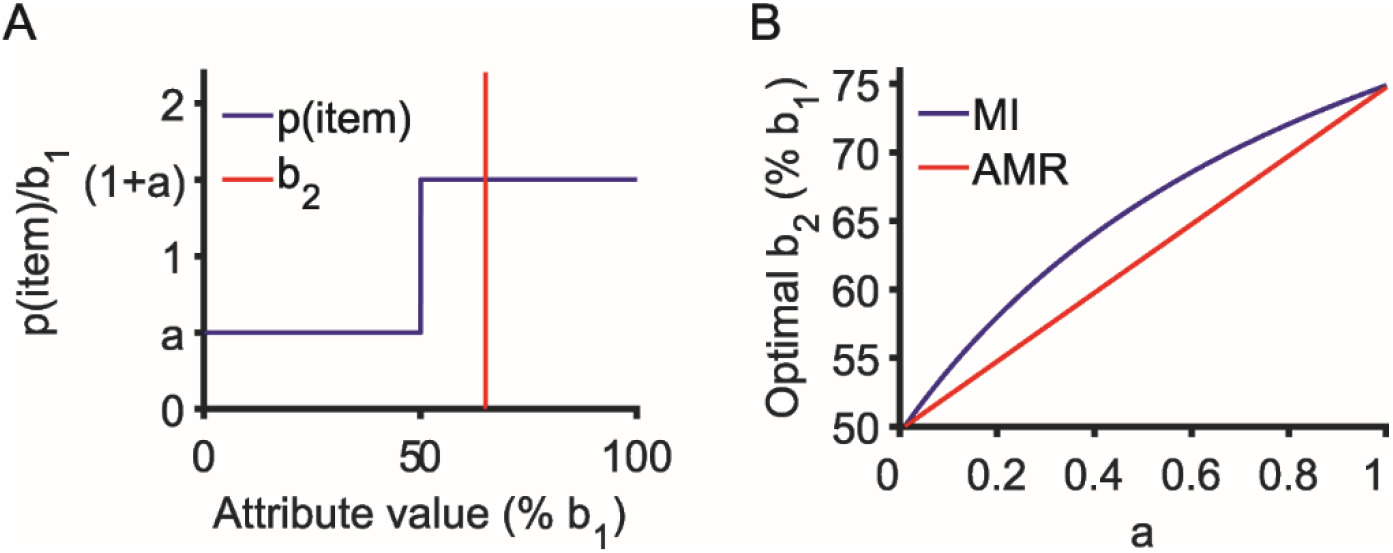
Item distribution and optimal percept boundary placement in E2B. (A) Panel shows the general shape of the distribution of item probabilities across the span of the value-determining attribute; panel also shows *b*_2_, the attribute value at which item space is perceptually split. The exact shape of the distribution is determined by the value of *a;* similarly, the exact position of *b*_2_ depends on *a* and on whether AMR or MI is being maximized. (B) Panel shows the relationship between *a* and the AMR- and MI-maximizing values of *b*_2_.

What I find in these sets of environments is that the compression strategies that maximize AMR and MI are rarely identical, but sometimes similar. In E2A_β1_, as shown in Part 3 of S1 Appendix, AMR is maximized at 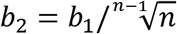 while MI is maximized at *b*_2_ = *b*_1_/2. In E2B, as shown in Part 3 of S1 Appendix and in Fig 1B, the AMR- and MI-maximizing values of *b*_2_ both increase strictly monotonically as a function of *a*, but do so at different rates, and are therefore equal only at *a* = 0 and *a* = 1. Together, these results suggest that, strictly speaking, perceptual optimality with respect to the MI objective is not evolutionarily valuable in its own right. This notion is not new: the lack of a direct link between perceptual MI maximization and organismal reward pursuit is a well-known shortcoming of the efficient coding model (1). That being said, if in some environments the AMR-maximizing compression strategy is a reasonably good approximation of both the evolutionarily optimal compression strategy and the MI-maximizing compression strategy, then in at least these environments, the predictions of the efficient coding model can perhaps be recapitulated within the optimal-fitness framework.

In order to better understand when AMR optimality is well-approximated by MI optimality, I first look to some environments in which it’s not: those in E2A_β1_ in which *n* is high. What is notable about the AMR-maximizing *b*_2_ value 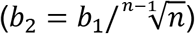 is that, as *n* increases, fit organisms will classify items as valuable (i.e., map them to the preferred percept) with increasing selectivity, and concomitantly lose their ability to differentiate amongst their environment’s less valuable items. Why is this evolutionarily favorable? As seen in Eq S6 in Part 2.1 of S1 Appendix, as *n* increases, so too does the probability that, for an arbitrarily chosen item value threshold, at least 1 item with value above that threshold will be presented. So, when these high-value items are likely to be present, natural selection favors the organisms that can differentiate these items from the rest.

The notion that AMR-maximizing compression strategies facilitate the selective identification of an environment’s most rewarding items suggests the following conjecture about the convergence of AMR- and MI-maximizing compression strategies. In some environments, the various decision scenarios, collectively, favor the selective identification of items that, as a set, exhibit a diversity of attribute values and populate many regions of item space. It is in these environments that the efficient coding model accurately approximates the AMR-maximizing compression strategy used by the environment’s organisms, because in these environments, maximizing differentiation amongst all of the environment’s stimuli is evolutionarily advantageous.

I illustrate this conjecture’s merit by optimizing compression in several environments within E2C and in E2D. In E2C, *β* = 2, *n* =10, and the values of the reward-determining attribute span a linear range. The specific E2C environments examined in this study, E2C.1, E2C.2, and E2C.3; are respectively comprised of 1, 2, and 4 decision scenarios; in which item reward values are maximized at attribute values (0), (0, *b*_1_), and (0, .15 * *b*_1_, .85 * *b*_1_, *b*_1_). The optimal compression strategies within these environments are derived in Part 5 of S1 Appendix and depicted in Fig 2. In E2C.1, the AMR-maximizing compression strategy excels at selectively identifying items with attribute value close to *b*_1_, and deviates significantly from the MI-maximizing compression strategy. Meanwhile, as conjectured, the AMR- and MI-maximizing compression strategies are similar in E2C.2, and even more so in E2C.3, because both environments favor selective identification of items at both ends of the item space, and because E2C.3 also favors selective identification of items within the item space’s interior. In E2D, the values of the reward-determining attribute span a radial range, there is a one-to-one correspondence between choice scenarios and attribute values (i.e., for every attribute value, there exists a scenario in which that attribute value maximizes reward), and the individual environments within the set are denoted by the parameter pair (*β, n*). In Part 6 of S1 Appendix, I demonstrate that within all E2D environments, the AMR- and MI-maximizing compression strategies are equivalent. This is notable because in the environments studied previously, high values of *n* conspicuously resulted in deviation from the MI-maximizing compression strategy in favor of the selective identification of the environment’s most valuable items; here, as conjectured, because all items are equally valuable, none are specifically targeted for selective identification when *n* is high.

**Fig 2.**
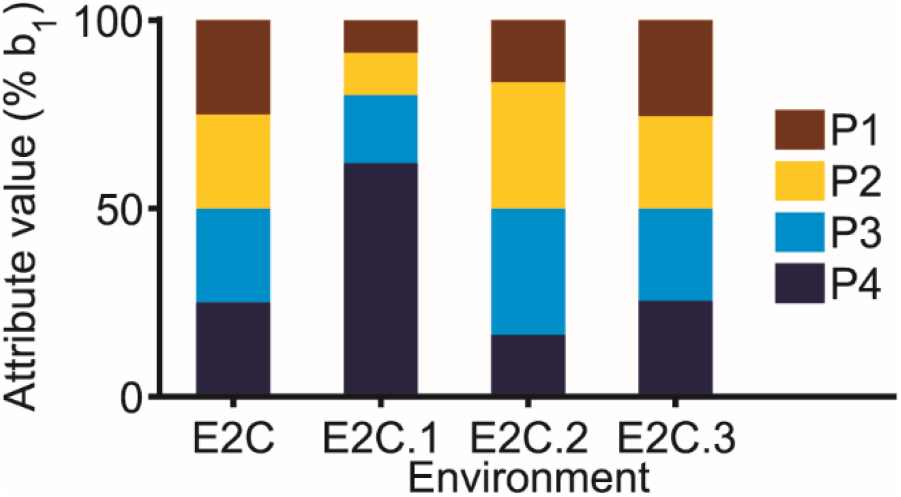
AMR-maximizing compression strategies in environments E2C.1, E2C.2, and E2C.3. Color-coded item-percept mappings depict the MI-maximizing compression strategy in all E2C environments, and the AMR-maximizing compression strategies in the environments indicated.

### Selection for deviation from information-theoretic optimality

E3 differs from the sets of environments considered previously in two significant ways. First, an organism’s fitness is based on reward accumulated over a finite lifespan (lifespan duration is measured in decisions), relative to other members of a fixed-sized population. More specifically: after every lifespan, the population undergoes a generational membership refresh during which the worst-performing accumulators leave no offspring. Second, environments within E3 occasionally present their inhabitant organisms with high-stakes (HS) decision scenarios. The organisms, for their part, use either an AMR-maximizing compression strategy or a bet-hedging (BH) compression strategy. The latter sacrifices optimal average-case performance in order to inform the avoidance of poor performance during HS scenarios (28, 35). Crucially, E3 contains environments in which lifespans are short, and thus, on account of unlucky AMR-maximizing organisms encountering HS scenarios and performing poorly, reproductive failure (and eventual extinction) is more likely for AMR-maximizing organisms than for BH organisms.

This study compares the fitness of these compression strategies within two subsets of E3. In both subsets, item probabilities are uniform over item space, and *n* = 2. In E3A, *β* = 1, and item reward value is a function of one attribute, as shown in Fig 3A. The AMR-maximizing placement of *b*_2_ is derived in Part 7 of S1 Appendix. In E3B, *β* = 2, there are two decision scenarios, and an item’s reward value in scenario 1 and scenario 2 is respectively equal to its attribute 1 and attribute 2 value. Notably, although *b*_1,1_ ≪ *b*_2,1_, scenario 2 is a rare occurrence, and thus, as shown in Part 8 of S1 Appendix, AMR maximization dictates the allocation of both transmitted bits to the differentiation of items along the span of attribute 1. Fig 3B shows the item-percept mappings of E3B’s AMR-maximizing and BH compression strategies.

**Fig 3.**
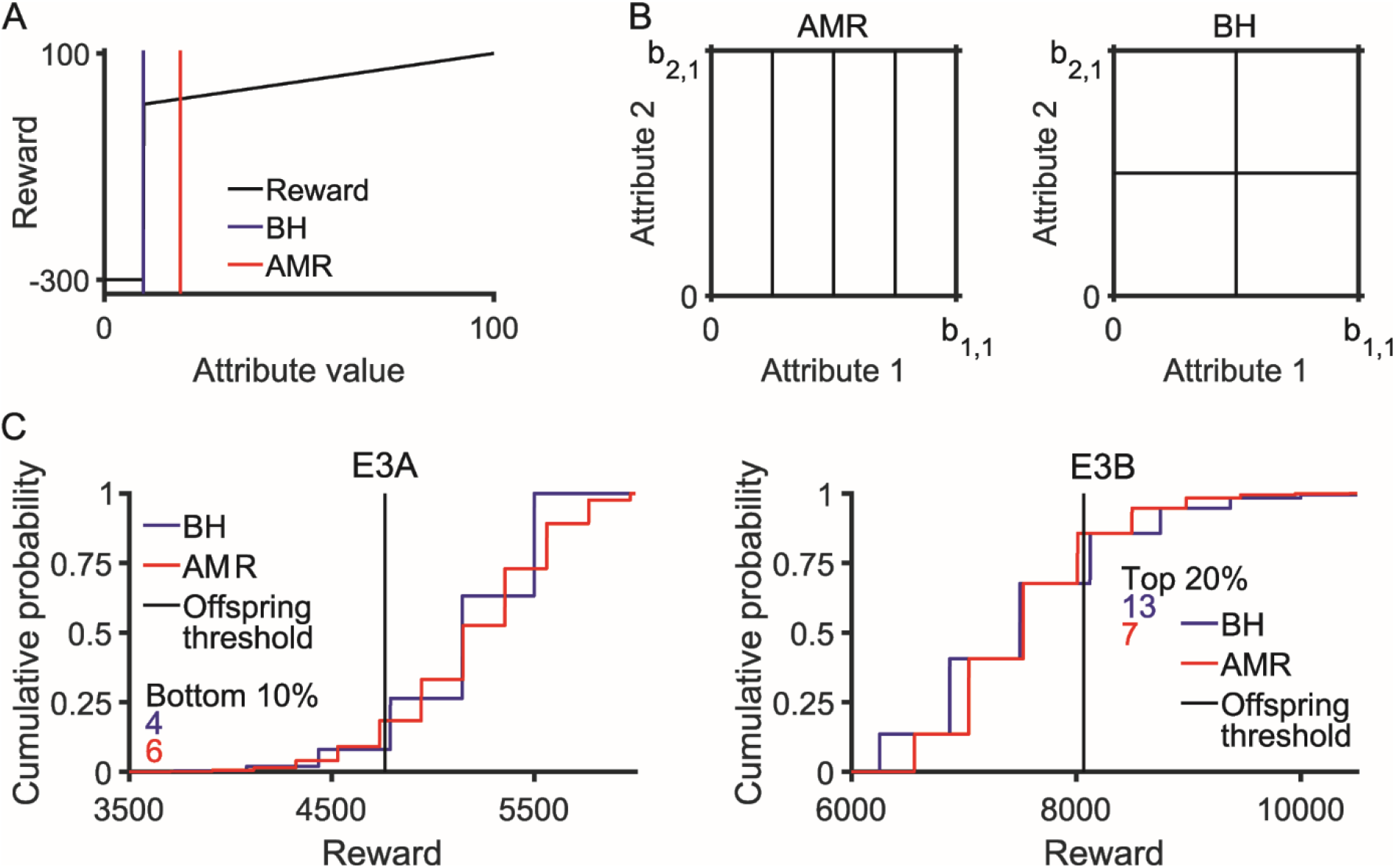
Compression strategy competition: AMR maximization vs. BH. (A) Relationship between attribute value and reward in E3A, along with the percept boundaries (*b*_2_) used by the AMR-maximizing and BH compression strategies. (B) E3B’s AMR-maximizing and BH item-percept mappings. (C) Probability distributions of lifetime reward accumulation, using the AMR-maximizing and BH compression strategies, in E3A and E3B. If an organism’s accumulated reward falls below the offspring threshold, that organism will not contribute to the subsequent generation. In populations with 50 AMR-maximizing and 50 BH organisms, the numbers of AMR-maximizing and BH organisms expected to be in E3A’s bottom 10% of reward accumulators, and in E3B’s top 20% of reward accumulators, are as indicated on each environment’s respective chart.

In both E3A and E3B, simulations of compression strategy competition mostly result in the population-wide fixation of AMR maximization; but, under certain conditions, BH prevails. As expected, in both E3A (Table 2) and E3B (Tables 3 and 4), BH fixation is favored when lifetimes are short. Meanwhile, the major difference between these environments is that in E3A, BH fixation is observed only when the population’s reproductive percentage is high (90%), while in E3B, BH fixation is observed only when the population’s reproductive percentage is low (20%). Fig 3C illustrates the genesis of this difference. In E3A, because AMR-maximizing organisms are less able to avoid the environment’s worst outcomes, they are more likely to fall into the population’s unreproductive 10%. In E3B, when only the most exceptional reward accumulators reproduce, selection favors BH organisms because they are more likely to obtain exceptionally large rewards during their encounters with HS scenarios. Interestingly, this E3B result is congruent with findings that high-variability strategies, i.e. risk-taking, can be advantageous in competitions that reward high relative standing (36, 37).

**Table 2.** Simulations of competition between AMR maximization and BH in E3A.

| **Lifetime**<br>**Decisions** | **BH, initial**<br>**prevalence** | **BH**<br>**fixations<sup>1</sup>** | **p-**<br>**value<sup>2</sup>** |
| --- | --- | --- | --- |
| 50 | 20% | 10 | .0020 |
| 50 | 50% | 10 | .0020 |
| 50 | 80% | 10 | .0020 |
| 1000 | 20% | 0 | .0020 |
| 1000 | 50% | 0 | .0020 |
| 1000 | 80% | 0 | .0020 |

<sup>1</sup>Each condition is simulated 10 times
<sup>2</sup>Binomial test, two-sided, n=10

**Table 3.** Simulations of competition between AMR maximization and BH in E3B.

| **Condition** | **Lifetime**<br>**Decisions** | **p(HS)<sup>1</sup>** | **BH, initial**<br>**prevalence** | **BH**<br>**fixations<sup>2</sup>** |
| --- | --- | --- | --- | --- |
| A | 2000 | .02 | 80% | 6 |
| B | 2000 | .015 | 80% | 0 |
| C | 2000 | .02 | 50% | 0 |
| D | 100 | .015 | 80% | 7 |
| E | 100 | .015 | 50% | 1 |
| F | 100 | .02 | 50% | 9 |
| G | 100 | .02 | 20% | 1 |

<sup>1</sup>Probability of HS scenario
<sup>2</sup>Each condition is simulated 10 times

**Table 4.** Comparisons of BH fixation probability in E3B.

| Conditions Compared<sup>1</sup> | Parameter difference | p-value<sup>2</sup> |
| --- | --- | --- |
| A vs. B | p(HS): .02 vs. .015 | 0.0108 |
| A vs. C | BH, initial prevalence: 80% vs. 50% | 0.0108 |
| D vs. B | Lifetime decisions: 100 vs. 2000 | 0.0031 |
| F vs. C | Lifetime decisions: 100 vs. 2000 | 0.0001 |
| D vs. E | BH, initial prevalence: 80% vs. 50% | 0.0198 |
| F vs. E | p(HS): .02 vs. .015 | 0.0011 |
| F vs. G | BH, initial prevalence: 50% vs. 20% | 0.0011 |

<sup>1</sup>Conditions as in Table 3, listed in descending order of BH fixation probability
<sup>2</sup>Fisher's exact test, two-sided, n=20

E4 is another set of environments in which the AMR-maximizing compression strategy is sometimes less fit than the BH compression strategy. In E4, item utilities are not fixed in evolutionary memory; instead, they are experienced and learned. Specifically, 100 items are randomly mapped to one of two percepts, each item’s reward value is sampled from a normal distribution N(μ,σ), and the organism then establishes a percept preference ordering by learning each percept’s expected value. It is assumed that percept values are learned quickly; and that a compression strategy’s fitness, aggregated over the population of organisms in a generation, is linearly proportional to either its single-decision AMR or its single-decision geometric mean reward (GMR).

As illustrated in Fig 4A, in the E4 environments in which item values are drawn from N(1,1) and *n* ∈ [9,10], the AMR-maximizing compression strategy’s pursuit of large rewards is fittest when fitness is based on AMR, while the BH compression strategy’s pursuit of steady rewards is fittest when fitness is based on GMR. Additional insight can be gained from examining why the AMR-maximizing compression strategy fails to produce the BH compression strategy’s steady rewards. Conceptually, the BH compression strategy potentially gets a reward sizably greater than μ from both percepts, while the AMR-maximizing compression strategy potentially gets a reward sizably greater than μ from only one percept, and thus, the latter’s GMR is lower because it’s pursuit of reward is undiversified (38). More specifically, the BH compression strategy maps 50 items to each percept, and thus, whichever percept is the most-preferred will deliver a reward that is somewhat greater than μ. Meanwhile, the AMR-maximizing compression strategy maps 90 items to one percept and 10 to the other. The 10-item percept has high variance, and thus confers the potential to provide an expected reward that is appreciably higher than μ. However, the 90-item percept is very similar to the environment’s total set of 100 items; so, when the 90-item percept is the preferred percept, the expected reward value of its items is only marginally higher than μ. The expected reward distributions of the 10-, 50-, and 90-item percepts, when they are the preferred percept, are shown in Fig 4B.

**Fig 4.**
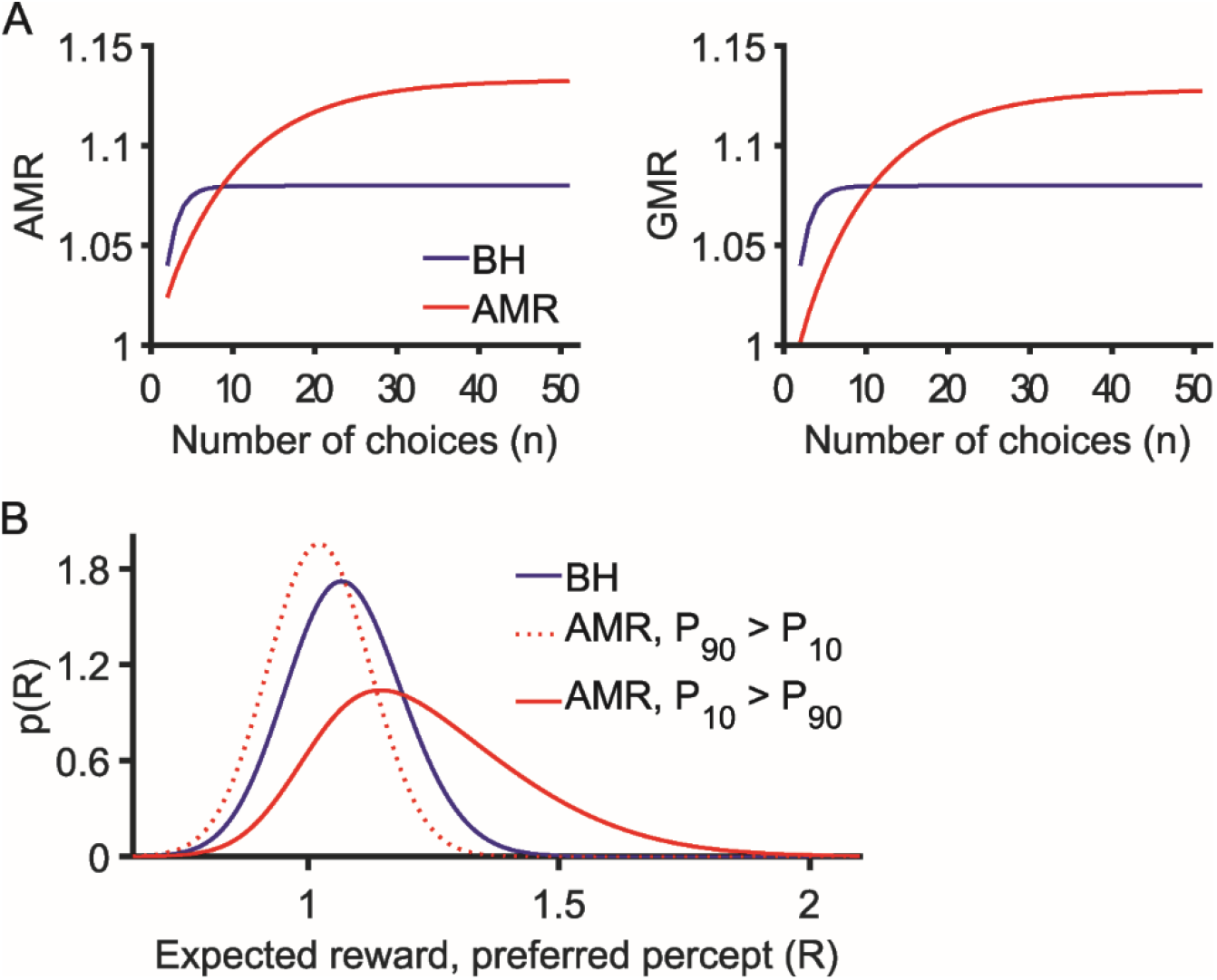
Optimizing the mapping of unknown-value items to percepts whose reward values are learnable. (A) AMR and GMR single-decision reward values obtained using a 1-bit perceptual system to distinguish amongst 100 items, the values of which are drawn independently from N(1,1). (B) Distributions of the expected value of items from the 50-item (BH), 10-item (AMR), and 90-item (AMR) percepts, when those percepts are the preferred percept.

Together, these findings from E3 and E4 demonstrate that natural selection can potentially favor compression strategies that are not predicted by the reward maximization model.

## Discussion

The optimal-fitness framework is a synthesis of various features of previously developed, and empirically well-supported, models of perceptual compression. From the matched filters model, it incorporates the premise that naturally-occurring perceptual systems are well-suited for the pursuit of evolutionary objectives, and the emphasis on the specific good decisions facilitated by a particular perceptual system. From information-theoretic models, it incorporates the notions that perception entails compression, and that compression must be optimized with respect to a particular objective. Taken together, these ideas result in a framework that entails 1) using compressed stimulus representations to 2) optimize reward 3) aggregated over the entire set of decisions potentially faced by the organism in its environment.

In the present study, the major conceptual advance enabled by the optimal-fitness framework is the reconciliation of the models that inspired it. Until now, there was no direct correspondence between the predictions of the efficient coding model and those of the reward maximization model, or between either information-theoretic model and the matched filters model. The results derived here, in E1 and E2, demonstrate correspondence between the models by expressing them in a shared mathematical language, and by delineating conditions under which a perceptual system’s properties can be simultaneously congruent with the matched filters model, the reward maximization model, and perhaps also the efficient coding model.

This study’s clarification of the relationship between these models, beyond being conceptually satisfying, has significant implications for the interpretation of empirical findings. Firstly, the deviations between AMR and MI maximization in Fig 1b and Fig 2, and the occasional selection for the BH compression strategy in E3 and E4, together serve to clarify that neither the AMR nor MI objective is always equivalent to fitness, which is ultimately the objective optimized by natural selection. Instead, recognizing that all models are wrong, but that some are useful, we see that the extant information-theoretic models are most useful when regarded as approximations of the compression strategies used by certain sensory systems. To understand these models as approximations is to be able to reconcile them with, for example, the existence of sensory systems that exhibit all of the following: efficient encoding of most naturalistic stimuli, reward-driven stimulus discrimination plasticity (21, 24-26), and adaptations that serve the processing of stimuli that inform essential survival actions (17-20).

Secondly, this study also highlights the existence of, and makes a preliminary attempt to address, the question of: under what conditions are these information-theoretic models most useful? In the absence of a theoretical framework for circumscribing the domain of application of these models, there is a risk that observed deviations from their predictions will be blithely regarded as reminders that these models are approximations, rather than as potentially-contradictory evidence. If the observations that would lead to the latter interpretation are not sufficiently well-defined, then these models fail to qualify as falsifiable!

In coming decades, it is likely that more work will be done, both experimentally and theoretically, to refine our understanding of the proper domain of application of these information-theoretic models. For time being, the results obtained in E1 and E3 suggest that the reward maximization model is more applicable to long-lifespan vs. short-lifespan organisms. Also, the results obtained in E2C and E2D suggest that the efficient coding model is most applicable to long-lived organisms that exploit the affordances associated with, and therefore benefit from selectively recognizing, a wide variety of stimuli. This latter finding is intriguingly congruent with empirical observations of efficient coding in the perceptual systems of behaviorally-complex vertebrates (defined here as those that interact with their environment in a great multitude of different ways). Moreover, the stimulus features that are efficiently encoded are often “low-level” features (such as edge orientation) (6, 33). Intuitively, it is easy to embrace the notion that humans and other long-lived behaviorally-complex vertebrates do it fact benefit from exploiting the affordances associated with a wide variety of stimuli and low-level stimulus feature combinations.

The optimal-fitness framework is additionally useful for modeling perceptual compressions that are not predicted by information-theoretic models, and are instead well-adapted for the accomplishment of various evolutionarily objectives that deviate from AMR maximization. In this study, this use of the optimal-fitness framework was demonstrated theoretically, via the presentation of models of compressions that were favored by natural selection due to their accomplishment of goals such as risk avoidance (in E3A), opportunistic resource pursuit (in E3B), and low-variability resource acquisition (in E4). To illustrate just one of many potential empirical applications of the optimal-fitness framework, I turn to the mating behavior of zebra finches (*Taeniopygia guttata*). Behaviorally, female zebra finches are mostly monogamous, they generally reject male attempts to initiate extra-pair copulation, and they preferentially initiate extra-pair copulation with males whose beak color indicates exceptional fitness (39-43). Perceptually, female zebra finches are more adept at perceiving color differences within some intervals of the male beak color range than within others, which would suggest that the distinctions made within the high-acuity interval are evolutionarily important (43). If the distribution of male beak colors were to be characterized, one would perhaps find that the high-acuity interval corresponds to the median beak color, which is what the efficient coding model would predict. Alternatively, one might find that the high-acuity interval separates the least-fit males from the rest. Would this compression be favored by selection if female finches avoided coupling up with low-fitness mates? Alternatively, one might find that the high-acuity interval separates the fittest males from the rest. Would this compression be favored by selection if there was significant benefit to engaging in extra-pair copulation with only the very fittest males, and could that benefit materialize even in light of the fact that extra-pair copulation is relatively rare? The optimal-fitness framework, which is able to express all of these behavioral strategies, along with an empirically-informed compression strategy and beak color distribution, could certainly be used to evaluate these questions. Moreover, if the construction of an environment that favors a particular ensemble of compression and behavioral strategies were to require the supposition of certain environmental features, the hypothesized existence of those features would likely be amenable to empirical verification.

Looking forward, there is certainly potential for developing, and gaining additional insight about the natural world from, more sophisticated versions of the optimal-fitness framework. Perhaps the most fruitful avenue for enhancing the current framework would be the addition of a time dimension. If, for example, a compression strategy were able to change across time, then it would be possible to model reward-driven stimulus discrimination plasticity. That, in turn, would enable the modeling of the above-mentioned hybrid compression strategies that mostly conform to efficient coding, but also exhibit sharpened tuning to a handful of exceptionally-rewarding stimuli. Taking time into account would also be useful for modeling explore-exploit decision environments in which decisions to exploit are followed by a fixed-time refractory period with no decisions presented at all, or in which decisions are only presented during certain time intervals.

In conclusion, the optimal-fitness framework has significant potential to continue to inform our understanding of the natural world. Already, in the present study, it has enabled the reformulation of the major extant perceptual compression models in a common mathematical language, demonstrated the potential overlap between their predictions, and addressed major outstanding questions about the breadth of applicability of information-theoretic models. Furthermore, it has been shown to straightforwardly facilitate the modeling of perceptual systems that are both evolutionarily fit and optimized for the pursuit of objectives other than AMR.

## Materials and Methods

### Compression strategy competition, E3

The procedure described here is used in both E3A and E3B. Three populations are simulated in parallel. The size of each is fixed at 100 individuals, which use either the AMR maximization or BH compression strategy. At simulation initialization, the population prevalence of AMR maximization is either 20%, 50%, or 80%; all three populations are initialized equivalently. Then, within each population, a lifetime of binary (that is, *n* = 2) item decisions is simulated for each member (the kth member of each population faces the same life decisions), the members are ranked by their lifetime’s accumulated reward, and the membership of the next generation is generated according to rules specific to each population. Lifetime simulation and population refresh procedures repeat until each population converges to fixation of one of the two compression strategies.

Binary decisions are simulated as follows: first, two items are drawn at random from the environment’s available items; then, either the more preferable item is chosen if the organism distinguishes the two items, or else the decision is made randomly.

The generational refresh procedures for the first and second populations are simple: the bottom 80% and 50% (respectively) of the reward accumulators are eliminated, and those remaining are copied until the population size is restored to 100. In the third population, the bottom 10% is eliminated, and each member thereof is replaced with an individual whose probability of using each compression strategy is proportional to the compression strategy’s prevalence in the top 90%.

In E3B, items are sampled from the item space in Fig 3B, with b1,1=100 and b2,1=1100. Decisions contingent upon attribute 2 are deemed high-stakes (HS), and arise with probability p(HS). In these simulations, p(HS) = .02 or .015; thus, p(HS)<.022, and therefore, AMR is maximized (as shown in S1 Appendix, Part 8) by allocating both transmitted bits to the differentiation of items along the span of attribute 1.

### Expected reward distributions, E3

The points on each cumulative probability distribution in Fig 2C represent both a probability and an accumulated reward amount. When computing the probability, it is assumed that the lifetime number of HS decisions faced by an organism is sampled from a Poisson distribution. When computing the accumulated reward, it is assumed that organisms obtain the expected reward from all decisions, where the expected reward depends on the compression strategy and on whether the organism is facing a HS or non-HS (nHS) decision scenario (this approximation is more accurate for nHS decisions, because in a lifetime there are many more of those). Thus, each distribution is determined by E(nHS), E(HS), and p(HS); the values of these parameters, for each distribution, are given in Table 5.

**Table 5.** Expected reward distribution parameters.

| Environment | Compression Strategy | $E(nHS)$ | $E(HS)$ | $p(HS)$ |
| --- | --- | --- | --- | --- |
| E3A | AMR | 59.75 | -146.66 | .038 |
|  | BH | 55 | -300 | .01 |
| E3B | AMR | 65.6 | 550 | .02 |
|  | BH | 62.5 | 687.5 | .02 |

The above parameters are determined as follows. In E3A, for both compression strategies, p(HS)=(b_2_/b_1_)^2^ and E(nHS)=(b_1_+b_2_)/2; for BH and AMR, E(HS) is respectively −300 and −146.66 (the latter is computed in S1 Appendix, Part 7). In E3B, p(HS) is manually set to .02, which is one of the values used in simulations, while the expected utilities are informed by the results in S1 Appendix, Part 8.

### Expected reward computation, E4

Because percepts are composed of items whose reward values are drawn from normal distributions, the distributions of mean percept values are also normal. Also, once percept values are learned, the organism will preferentially select items mapped to higher-values percepts; a percept’s expected value, therefore, is related to its position in the percept preference order. If it is further specified that the organism maps items to *N* percepts, and that a particular sampling of item reward values results in percept 1 being the highest-value percept, then the expected value of percept 1, *E*(*P*_1_), can be expressed as

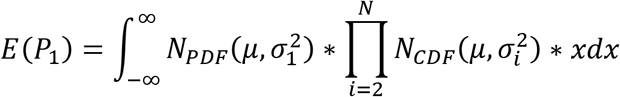

Alternatively, if *E*(*P*_1_) < *E*(*P*_2_) and *E*(*P*_1_) > *E*(*P_i_*) ∀ *i* >2, then

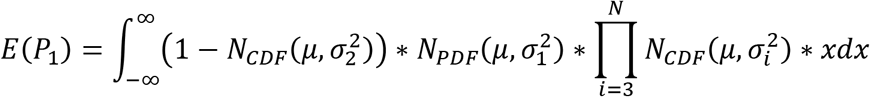

In this study, the expressions for all expected percept values across all preference orderings are constructed similarly. Within these expressions, the σ-value associated with each *N_CDF_* and *N_PDF_* is computed using standard methods: for example, for a 25-item percept with item reward values drawn from N(0,1), σ=1/5. Once the expressions are defined, they are evaluated by numerical integration: the results in Fig 4A (4B) are obtained by integrating over [−5,5] ([.5,2.25]) with dx=.001.

Upon computing the ordering-specific *E*(*P_i_*), it becomes possible to combine them to compute the mean reward obtained from a particular mapping of 100 items to the available percepts. Specifically, a mapping’s respective AMR and GMR are 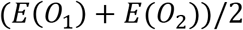 and 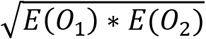, where *O_j_* denotes the ordering in which *j* is the preferred percept. Defining *E*(*P_j,i_*) as the expected value of percept *i* in ordering *j*, defining *S_i_* as the size (in items) of percept *i*, and recalling that *n* items are presented per decision,

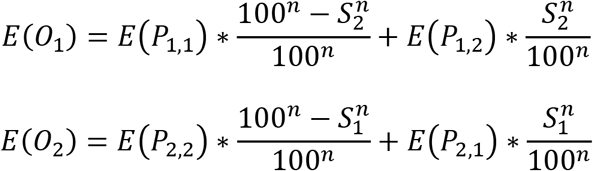

## S1 Appendix. Mathematically-derived results

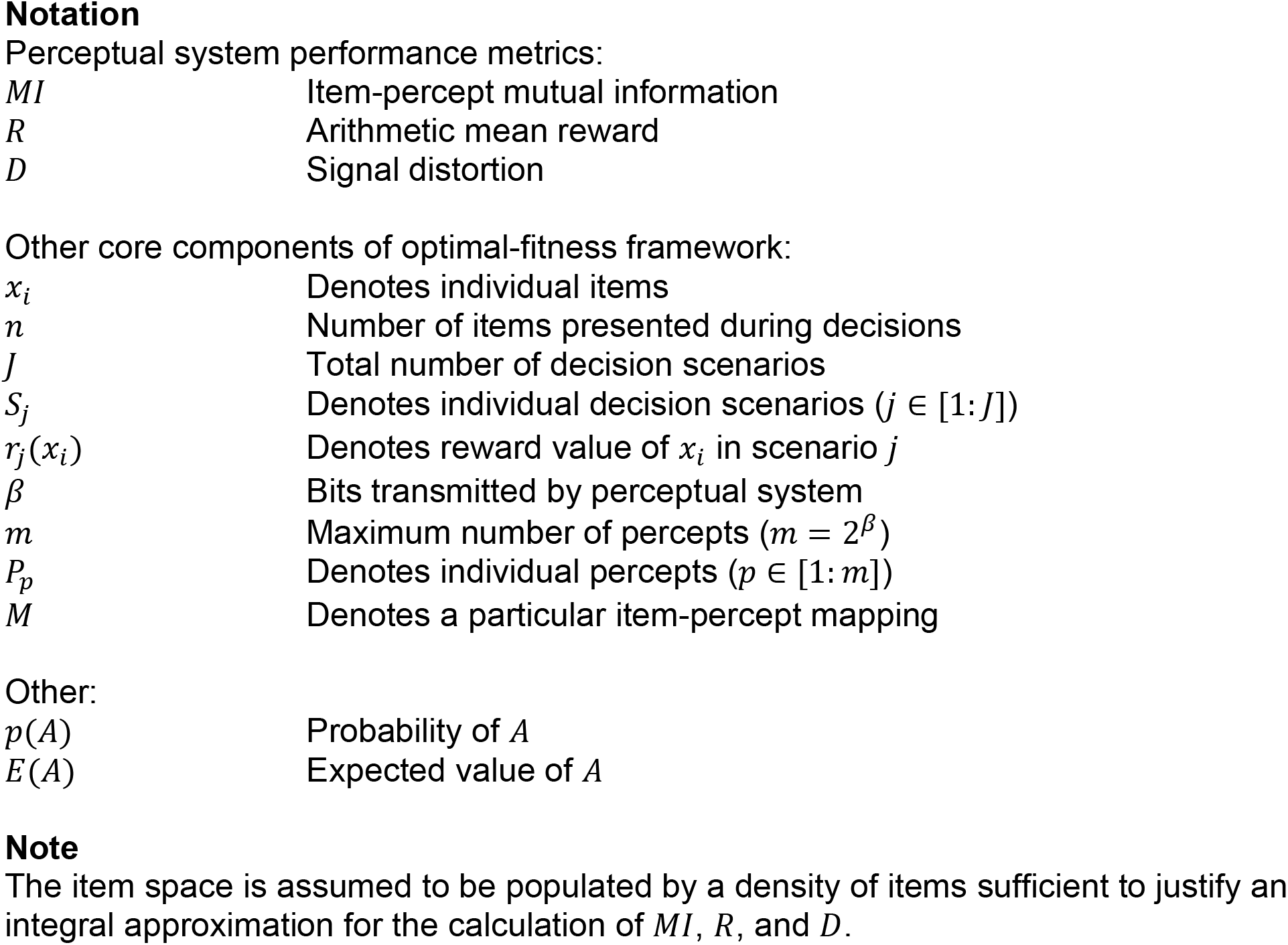

|  |  |
| --- | --- |
| $MI$ | Item-percept mutual information |
| $R$ | Arithmetic mean reward |
| $D$ | Signal distortion |
| $x_i$ | Denotes individual items |
| $n$ | Number of items presented during decisions |
| $J$ | Total number of decision scenarios |
| $S_j$ | Denotes individual decision scenarios ($j \in [1:J]$) |
| $r_j(x_i)$ | Denotes reward value of $x_i$ in scenario $j$ |
| $\beta$ | Bits transmitted by perceptual system |
| $m$ | Maximum number of percepts ($m = 2^\beta$) |
| $P_p$ | Denotes individual percepts ($p \in [1:m]$) |
| $M$ | Denotes a particular item-percept mapping |
| Other: |  |
| $p(A)$ | Probability of $A$ |
| $E(A)$ | Expected value of $A$ |

### Part 1: Maximizing *R* and minimizing *D* in E1

#### 1.1: *R* (noiseless item-percept mapping)

In E1, fitness is proportional to *R*. To optimize perception with respect to *R*, note that for a particular organism in a particular environment, the following variables are fixed: item probabilities over the attribute space, choice scenarios, and *m*. Thus, only *M* is subject to natural selection, which lets the perception optimization problem be expressed as 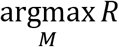, where 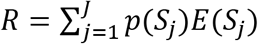 and

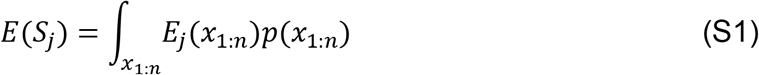

Conceptually, Eq S1 represents the sum of the expected reward obtained from all sets of *n* items, weighted by each set’s occurrence probability. Note that *x*_1:*n*_ is shorthand for *x*_1_, …, *x_n_*.

*M* affects *E_j_*(*x*_1:*n*_) as follows: when the items in a particular set *x*_1:*n*_ are mapped to their respective percepts, *K* (1 ≤ *K* ≤ *n*) items will be mapped to a percept *P_p_* that the organism deems preferable to the others. Thus, 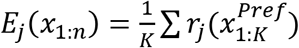, where 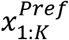 represents the items mapped to the preferred percept. Note that for some item sets, the preferred percept (*P_Pref_*) will be *P*_1_ for others, *P*_2_; similarly for all *P_p_*(*p* = 1:*m*).

#### 1.2: *D* (noiseless item-percept mapping), and equivalence between maximizing *R* and minimizing *D*

Optimizing *D* is fundamentally similar to optimizing *R*. In this case, the objective can be stated as 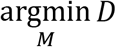, with 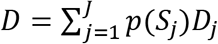 and

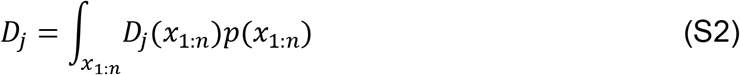

If distortion is defined with the cost function of reward forfeited as a result of compression, then *M* affects *D_j_*(*x*_1:*n*_) as follows. If compression resulted in no information loss, then 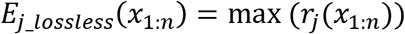. However 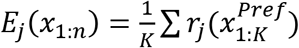. Thus, 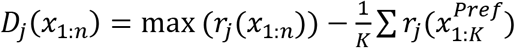, and

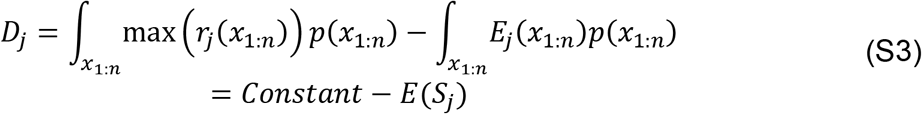

Thus, the *M* that maximizes *R* minimizes *D*.

#### 1.3: Compensating for noise

In the context of this study’s item-percept mapping framework, noise can be implemented as follows: with probability 1 – *δ* (*δ* small), an item is perceived as an instance of its target percept *P_T_*; otherwise, it is mapped to other percepts *p*(*p* ≠ *T*) with probability *δ_p_*(*δ* = ∑ *δ_p_*).

To express how item-percept mapping noise affects *R*, note that in Eq S1, noise would be expected to affect only the *E_j_*(*x*_1:*n*_) term. Denoting a particular noisy mapping of *x*_1:*n*_ as *M^Noi^*, and the set of all *M^Noi^* for a particular *x*_1:*n*_ as *MAP*, yields

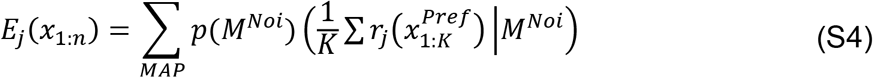

Here, the meaning of 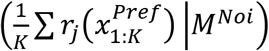 is that, for a particular *M^Noi^*, the expected reward is the average value of the *K* items mapped to the most-preferred percept presented by *M^Noi^*.

Clearly, there is potential for the noise-adjusted optimal mapping to differ from the *M* that is optimal when all *δ_p_* = 0. A concrete example of difference between these mappings is provided in Part 2.2, below.

### Part 2: Maximizing *R* in E2A

#### 2.1: Optimal *M* (noiseless item-percept mapping)

In E2A, item probabilities are uniform, and there is just one choice scenario. Thus, the expression of *R* is simplified: *p*(*S*_1_) = 1, *R* = *E*(*S*_1_), *p*(*x*_1:*n*_) is constant, and

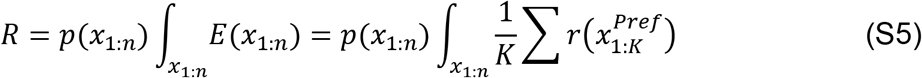

Next, note that in E2A, items are most valuable at attribute value *b*_1_, boundaries are indexed in order of proximity to *b*_1_, percepts are indexed according to their upper boundary, and therefore, percepts are indexed in order of their desirability. Thus, for a given item set *x*_1:*n*_, the probability that percept *p* will be *P_Pref_* is

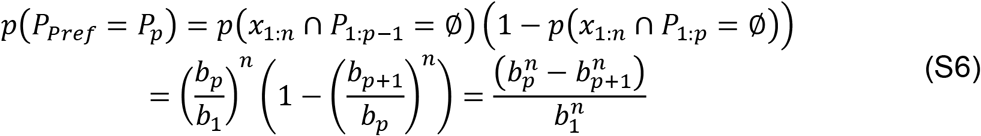

Next, note that when 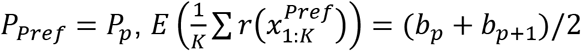. Thus,

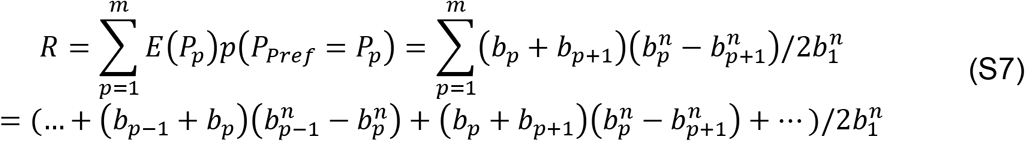

To determine optimal boundaries, set *dR/db_p_* = 0 and solve, which yields

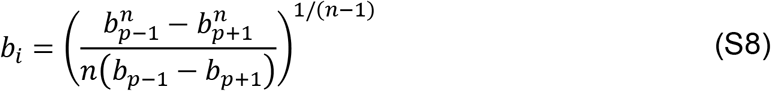

#### 2.2: Comparing optimal *M* in noise’s presence vs. absence

In E2A, with sufficiently low *β* and *n*, the noise-adjusted optimal mapping can be determined without undue difficulty. So, to illustrate how noise affects the optimization of *M*, I derive the optimal *M* when *β* = 1, *n* = 3, and *δ* is small (≤.05).

In the following expression for *R* (Eqs S9-S11), the four summed terms respectively represent the events of 3, 2, 1, and 0 items sampled from *P*_1_. Each of these sampling events is expressed as *p*(*event*) * *E*(*event*). *A*_1_ and *A*_2_ are formulated similarly, with the top (bottom) line of each corresponding to the organism’s choosing of an item sampled from *P*_1_(*P*_2_).

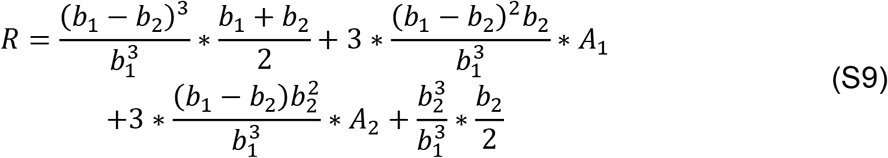

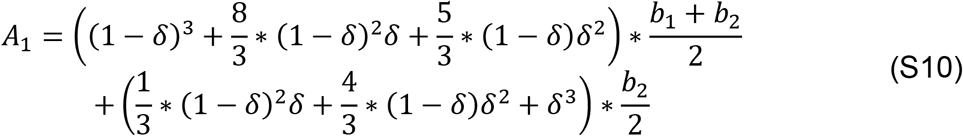

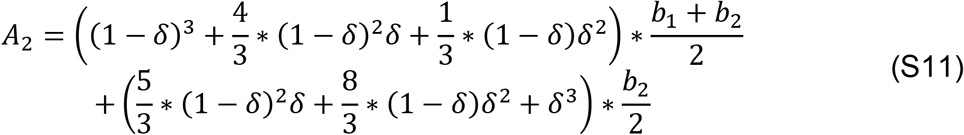

Setting *dR/db*_2_ = 0 yields

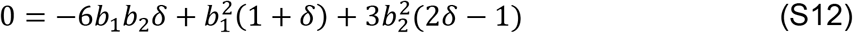

Note that when 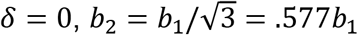, which is the result indicated in Eq S8. However, when *δ* = .05, *b*_2_ = .571*b*_1_.

### Part 3: Maximizing *MI* in E2A and *R* in E2A_β1_

#### 3.1: MI

If the identities of items spanning the attribute value range [0, *b_1_*] are guessed at random, their differential entropy is 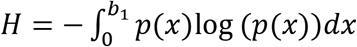. If, however, the items are mapped to one of *m* percepts, then their conditional entropy is

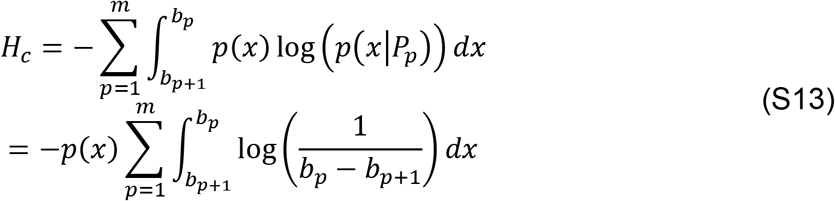

Note that *MI* = *H – H_c_* and *p*(*x*) = 1/*b*_1_. Setting *dH_c_/db_p_* = 0 and solving yields *b_p_* = (*b*_*p*–1_ + *b*_*p*+1_)/2. Because *b*_*m*+1_ = 0, *b*_*m*–1_ = 2 * *b_m_*. Also, if *b*_*p*+1_ = *n*x* and *b_p_* = (*n* +1) *\* x*, then *b_p–1_* = (*n* + 2) *\* x*. Thus, *b_m–n_* = (*n* + 1) *\* b_m_* and *b*_1_ = *m * b_m_*, and, as expected, *MI* is maximized when the boundaries are evenly spaced (by *b*_1_/*m*).

In E2A_β1_, *m* = 2 and therefore *b*_2_ = *b*_1_/2.

#### 3.2: R

Using Eq S8 and noting that *b*_3_ = 0 yields 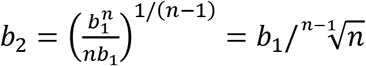.

### Part 4: Maximizing *R* and *MI* in E2B

In E2B, 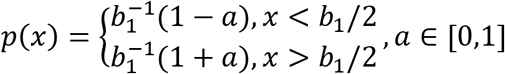

#### 4.1: R

Generalizing from Eq S6 and Eq S7, 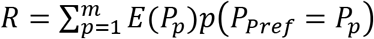. Thus,

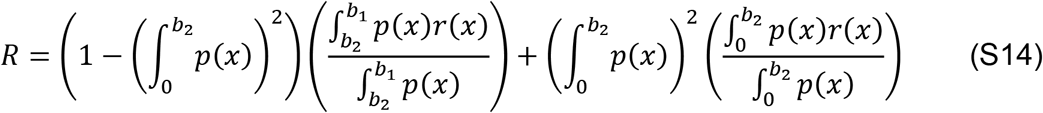

Because *p*(*x*) is constant in [*b*_2_, *b*_1_],

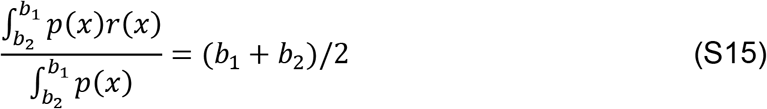

Also,

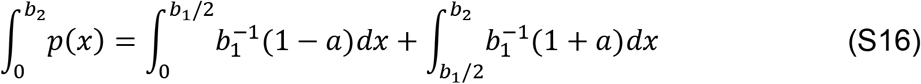

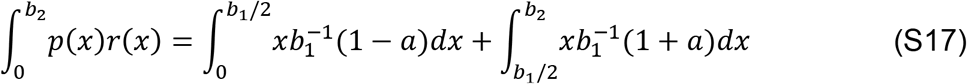

Setting *dR/db*_2_ = 0 and solving yields *b*_2_ = *b*_1_(2 + *a*)/4.

#### 4.2: MI

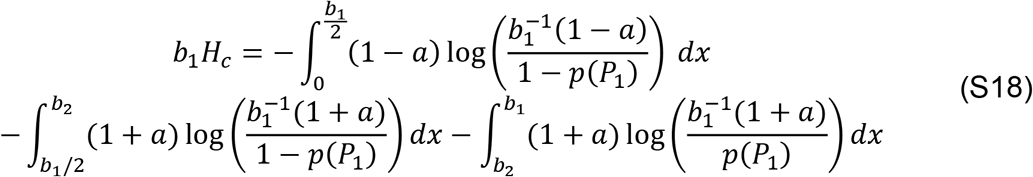

Letting *A*_1_ = *dp*(*P*_1_)/*db*_2_,

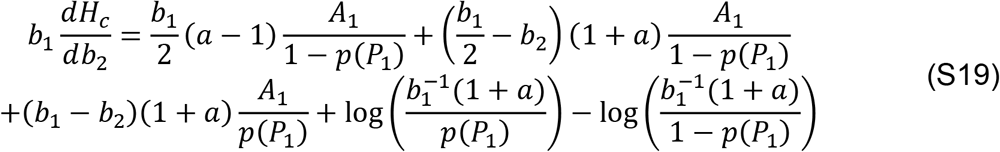

In Eqs S18 and S19, 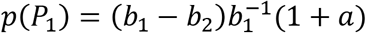. Setting *p*(*P*_1_) = .5 and solving yields *b*_2_ = *b*_1_(1 + 2*a*)/(2 + 2*a*). If *p*(*P*_1_) = .5, Eq S19 simplifies to

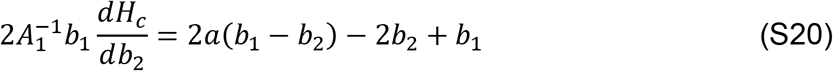

Setting *dH_c_/db*_2_ = 0 and solving yields *b*_2_ = *b*_1_(1 + 2*a*)/(2 + 2*a*). This *b*_2_ value matches that at which *p*(*P*_1_) = .5. Thus, when *b*_2_ assumes this value, *MI* is maximized.

### Part 5: Maximizing *MI* in E2C; and *R* in E2C.1, E2C.2, and E2C.3

#### 5.1: MI

Using the result of Part 3.1, because *m* = 4, *b*_4_ = *b*_1_/4, and thus, (*b*_2_, *b*_3_, *b*_4_) is optimized at *b*_1_ * (.75, .5, .25).

#### 5.2: E2C.1

Using Eq S8 and noting that 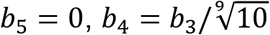, and

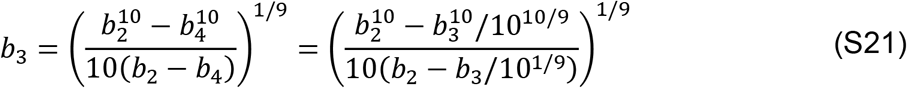

Numerical approximation yields *b*_3_ = .876*b*_2_. Similarly (letting *c* = .876),

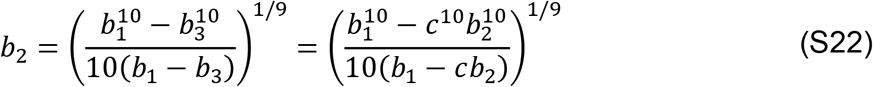

Numerical approximation yields *b*_2_ = .915*b*_1_.

Thus, (*b*_2_, *b*_3_, *b*_4_) is optimized at *b*_1_ * (.915, .801, .62).

#### 5.3: E2C.2

*R* = *E*(*S*_1_) + *E*(*S*_2_), and in *S*_1_ (*S*_2_), item reward is maximal at *b*_1_ (0). From Eq S7,

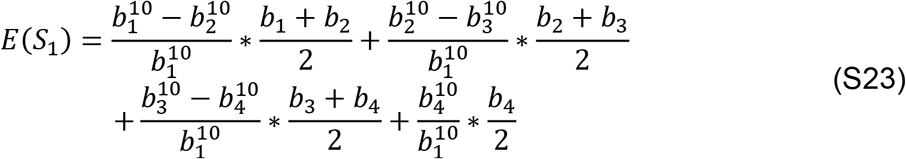

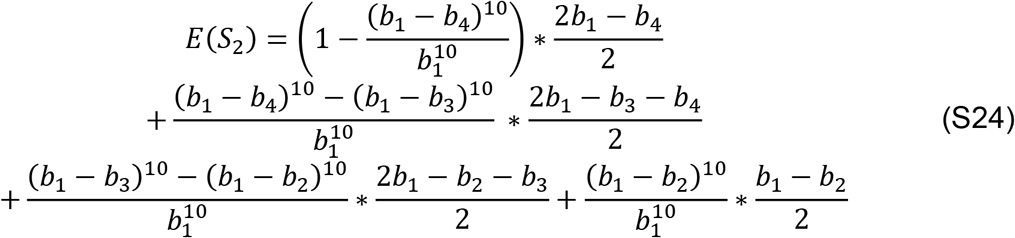

The search for optimal (*b*_2_, *b*_3_, *b*_4_) can begin by optimizing with respect to *b*_3_:

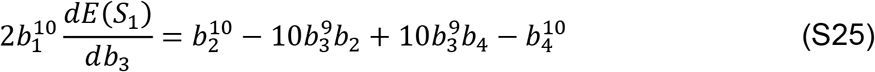

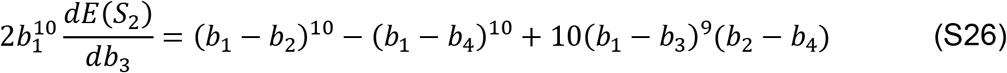

If it is assumed that *b*_4_ = *b*_1_ – *b*_2_, then setting *dR/db*_3_ = 0 and solving yields *b*_3_ = *b*_1_/2.

Substituting *b*_4_ = *b*_1_ – *b*_2_ and *b*_3_ = *b*_1_/2 into Eq S23 and Eq S24, it becomes clear that *E*(*S*_1_) = *E*(*S*_2_), which is as expected. Setting *dR/db*_2_ = 0 and numerically approximating yields *b*_2_ = .836*b*_1_; thus, (*b*_2_, *b*_3_, *b*_4_) is optimized at *b*_1_ * (.836, .5, .164).

#### 5.4: E2C.3

*R* = *R*_1_ + *R*_2_, where *R*_1_ = *E*(*S*_1_) + *E*(*S*_2_) and *R*_2_ = *E*(*S*_3_) + *E*(*S*_4_). To characterize *R*_2_, note, first, that in *S*_3_, in which item reward value peaks at an attribute value. 15*b*_1_ removed from attribute value corresponding to peak item reward value in *S*_1_, the most-preferred percept is the same as in *S*_1_. Next, note that for percepts 2, 3, and 4, *E*_3_(*P_p_*) = *E*_1_(*P_p_*) + .15*b*_1_ (recall that in *E_j_*, subscript *j* denotes the scenario). Finally, note that *S*_4_ can be described with reference to *S*_2_ in a manner analogous to the description of *S*_3_ with reference to *S*_1_.

Thus,

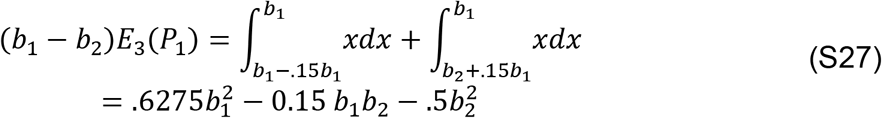

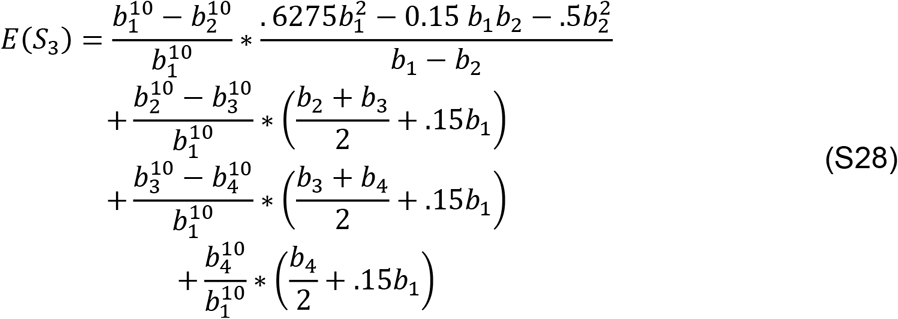

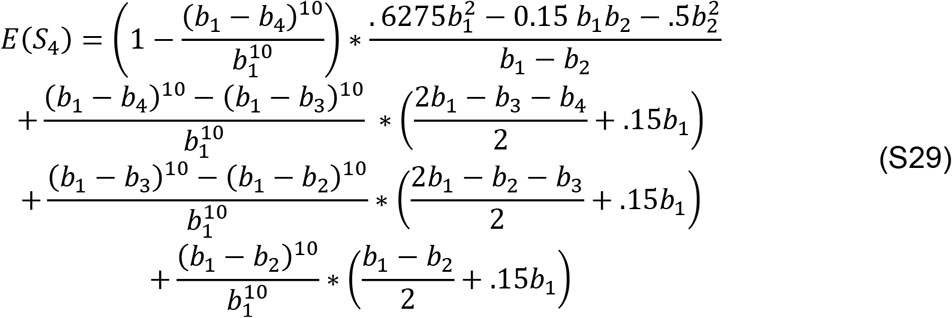

Optimizing Eq S28 and Eq S29 with respect to *b*_3_ yields Eqs S25 and Eq S26, respectively. So, if it is again assumed that *b*_4_ = *b*_1_ – *b*_2_, then setting *dR*_2_/*db*_3_ = 0 and solving yields *b*_3_ = *b*_1_/2. Thus,

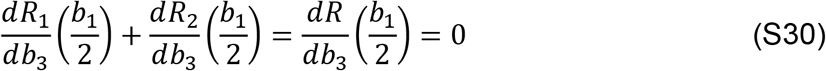

Substituting *b*_4_ = *b*_1_ – *b*_2_ and *b*_3_ = *b*_1_/2 into Eq S28 and Eq S29, it becomes clear that *E*(*S*_3_) = *E*(*S*_4_), which is as expected. Setting *dR/db*_2_ = 0 and numerically approximating yields *b*_2_ = .746*b*_1_; thus, (*b*_2_, *b*_3_, *b*_4_) is optimized at *b*_1_ * (.746, .5, .254).

### Part 6: Equivalence of compression strategies that optimize *R* and *MI* in E2D

#### 6.1: MI

Without loss of generality, an arbitrarily-chosen percept boundary within E2C can be labeled *b*_1_, and the rest, moving counterclockwise from *b*_1_, can be labeled *b*_2:*m*_. Next, the circle of E2C’s attribute values can be “linearized”, by “cutting” at *b*_1_ and relabeling the rightmost edge of *P_m_* as *b*_*m*+1_, without affecting item-percept *MI* because the linearization does not affect item-percept mappings. Now, because of the uniformity of item probabilities, the result of Part 3.1 is applicable here: evenly-spaced boundaries maximize *MI*.

#### 6.2: Paired decision scenarios

In an effectively unidimensional environment, decision scenarios *S*_1_ and *S*_2_ can be defined as “paired with respect to *b_i_*” if the following conditions are met:

1. *S*_1_ and *S*_2_ are equally probable.
2. Outside of the interval [*b*_*i*–1_, *b*_*i*+1_], percept boundaries are positioned such that, for every percept in *S*_1_, there exists a percept in *S*_2_ with equivalent expected value and cumulative item probability.
3. Item probabilities within [*b*_*i*–1_, *b*_*i*+1_] are uniform.
4. The relationship between item attribute value and scenario-specific reward value obeys 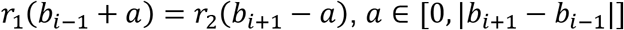.

If the above conditions are met, it can be shown that *b_i_* = (*b*_*i*–1_ + *b*_*i*+1_)/2 is an optimum of *E*(*S*_1_) + *E*(*S*_2_). To do so, it is helpful to denote as *P*_1_(*P*_2_) the percept bordered by *b*_*i*–1_ and *b_i_* (*b_i_* and *b*_*i*+1_), and, similarly to Eq S1 and Eq S7, denote in scenario *j* a percept’s expected value as *E_j_*(*P_p_*) and probability of being the most-preferred percept as *p_j_*(*P_p_*).

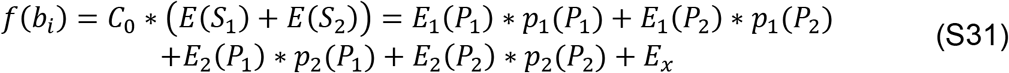

Here, 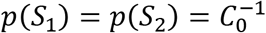 on account of condition (1), and *E_x_* refers to all terms in *f*(*b_i_*) with no direct dependence on *b_i_*. Differentiating with respect to *b_i_* yields

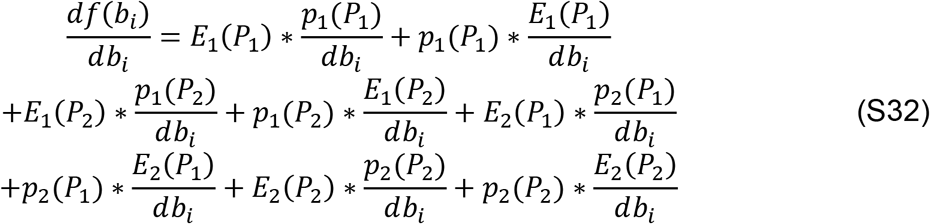

When *b_i_* = (*b*_*i*–1_ + *b*_*i*+1_)/2, then, on account of conditions (3) and (4), *E*_1_(*P*_1_) = *E*_2_(*P*_2_) = *C*_1_, *E*_1_(*P*_2_) = *E*_2_(*P*_1_) = *C*_2_, 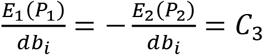, and 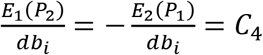.

On account of this result and conditions (2) and (3), *p*_1_(*P*_1_) = *p*_2_(*P*_2_) = *C*_5_, 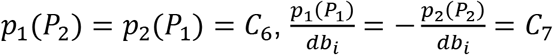, and 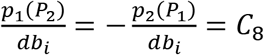. Thus,

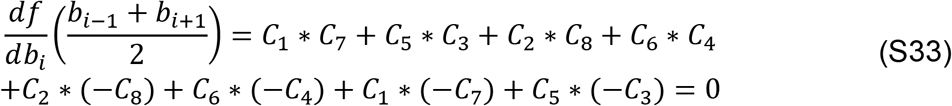

#### 6.3: R

Similarly to Part 6.1, and because of the environment’s symmetry, *b*_1_ can be placed arbitrarily, with the other boundaries, counterclockwise from *b*_1_, being labeled *b*_2:*m*_. Also, traversing the environment’s attribute value in the counterclockwise direction can be defined as positive angular motion. Consider now the decision scenarios in which reward peaks at attribute values *b*_2_ + *a* and *b_m_* – *a*, respectively labeled *S_a_* and *S_−a_*. If, for arbitrary *a* ∈ (*-π/m, π – π/m*), *S_a_* and *S_−a_* are paired with respect to *b*_1_, then

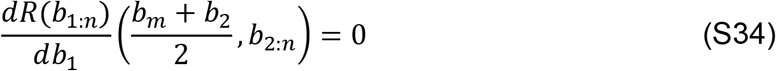

where

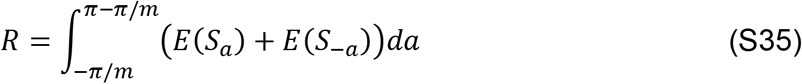

When *S_a_* and *S_−a_* are paired with respect to *b*_1_, |*b*_1_ – *b*_2_| = |*b*_1_ – *b_m_*| = 2*π/m*. If they are paired when 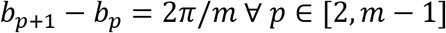, then *R* is optimized when percept boundaries are evenly spaced. Recalling now that any of the boundaries can be labeled *b*_1_, it follows that for any boundary *p*, if all other boundaries are spaced 2*π/m* apart, then 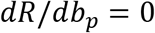 at 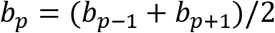. Thus, 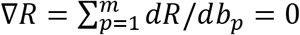 when 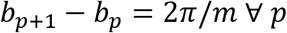, and *R* and *MI* are optimized by the same compression strategy.

It remains to be demonstrated that, for arbitrary *a* ∈ (-*π/m,π – π/m*), *S_a_* and *S_−a_* are paired under the following conditions: their respective reward peaks occur at attribute values *b*_2_ + *a* and *b_m_ – a*, and *b*_*p*+1_ – *b*_p_ = 2*π/m* ∀ *p ∈* [2, *m* – 1]. *S_a_* and *S_−a_* are paired if the conditions presented in Part 6.2 are met. Conditions 1 and 3 are met by virtue of the properties of E2D. Condition 4 is met as well: *r_a_*(*b*_2_) = *r_−a_*(*b_m_*) = *M – a* and *r_a_*(*b_m_*) = *r_−a_*(*b*_2_) = *M – a* – 4*π/m*. Finally, condition 2 is met because 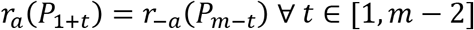.

### Part 7: Maximizing *R* in E3A

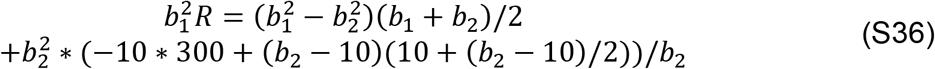

Setting *dR/db*_2_ = 0 and solving yields *b*_2_ = 19.5. A further useful result is expected value of the non-preferred percept when *b*_2_ = 19.5:

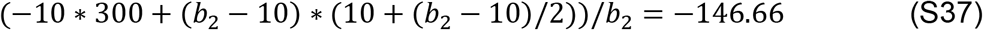

### Part 8: Maximizing *R* in E3B

In E3B, *R* = *E*(*S*_1_)*p*(*S*_1_) + *E*(*S*_2_)(1 – *p*(*S*_1_)). Note that maximizing *R* entails the consideration of three configurations: *R*_11_ and *R*_22_, in which both bits are respectively allocated to differentiating items along the span of attribute 1 and attribute 2; and *R*_12_, in which one bit is allocated to differentiating items along each attribute’s span. Also, for simplicity, I normalize *b*_1,1_ and *b*_2,1_ by *b*_1,1_; thus, *b*_1,1_ → 1 and *b*_2,1_ → 11.

In *R*_11_, *E*(*S*_2_) = 11/2. The optimal placement of *b*_1,2_, *b*_1,3_, and *b*_1,4_ (*b*_1,5_ = 0) can be determined by using Eq S8; because *n* = 2,

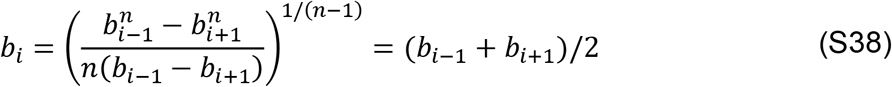

From Parts 3.1 and 5.1, we know what boundary placements maximize an objective of interest (in this case, *R* rather than *MI*) when *b_i_* = (*b*_*i*–1_ + *b*_*i*+1_)/2 and *m* = 4.

Accordingly, (*b*_1,2_, *b*_1,3_, *b*_1,4_) is optimized at (.75,.5,.25).

From Eq S7, the reward obtained using this boundary placement is

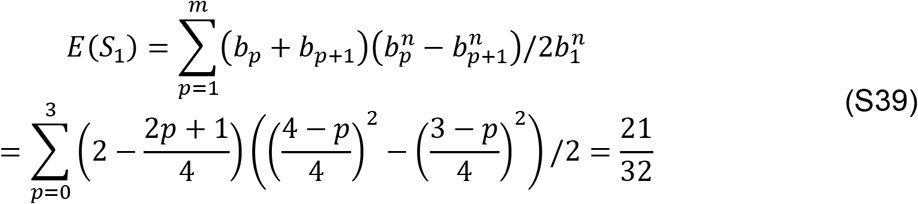

Thus, 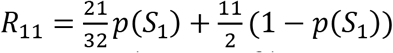.

Similarly, 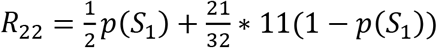.

Meanwhile, 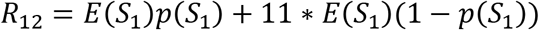.

In 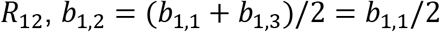.

Accordingly, 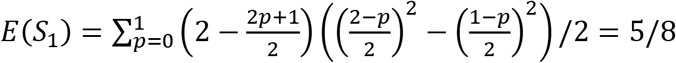.

Thus, *R*_22_, *R*_12_, and *R*_11_ respectively maximize R when *p*(*S*_1_) > .266, when .022 > *p*(*S*_1_) > .266, and when *p*(*S*_1_) < .022.

